# CurvoChip: a programmable dynamic curvature-on-chip platform for epithelial mechanobiology

**DOI:** 10.64898/2026.08.21.746327

**Authors:** Rémi Tranzer, Charlotte Rivière, Alejandro Ibarra, Marine Luciano, Sylvain Gabriele

## Abstract

Epithelial tissues continuously remodel their curvature during morphogenesis, homeostasis, regeneration, and disease, yet experimental access to time-varying curvature remains limited. Here, we introduce CurvoChip, a pneumatically actuated microsystem that reversibly deforms confluent epithelial monolayers cultured on a 20-µm elastic membrane into concave or convex geometries. The device operates either in a standard incubator or on a microscope stage and provides programmable control over pressure amplitude, direction, and cycling. Analytical scaling, finite-element simulations, and confocal profilometry establish predictable membrane deformation across the operating range, whereas cycling between −400 and +400 mbar for 120 cycles produces stable deflection without detectable drift or residual deformation. We further implement a three-dimensional surface-reconstruction and segmentation workflow to quantify cell and nuclear morphology on curved monolayers. Acute curvature induction produces a marked polarity-dependent response: convex deformation causes greater cell spreading and epithelial thinning than concave deformation, while nuclear projected area, thickness, and volume change in a direction- and position-dependent manner. These results show that epithelial architecture is sensitive not only to curvature magnitude but also to its orientation relative to the apico-basal axis. CurvoChip therefore provides an accessible platform for dissecting how epithelial tissues integrate dynamic geometric cues.

## Background

Curvature is a fundamental geometric feature of biological organization. The three-dimensional architecture of epithelial tissues shapes organ structure and function, from neural-tube folding to the curved alveolar surfaces of the lung (1, 2). In mechanically active organs such as the lung and intestine, epithelial geometry also fluctuates over time, coupling tissue deformation to biochemical signaling (3). Nevertheless, the amplitudes and characteristic timescales of the curvature changes experienced by epithelia in vivo remain poorly quantified. Most in vitro models rely on static or preformed curved substrates and therefore provide limited access to reversible or cyclic changes during live imaging (4, 5). They consequently capture only part of the dynamic geometry associated with breathing, peristalsis, regeneration, and morphogenetic remodeling.

The acquisition and maintenance of a three-dimensional epithelial shape require coordinated control of cell deformation, tissue stress, and luminal pressure. Transmural pressure has been exploited as a morphogenetic driver in systems in which weakly adherent MDCK monolayers form fluid-filled cavities on micropatterned substrates. Osmotically driven swelling of these cavities generates dome-shaped epithelia and has revealed that curvature can regulate stress anisotropy and cell alignment independently of tissue size (6). In such systems, however, curvature is principally an emergent outcome of cavity inflation rather than an externally prescribed variable that can be reversed repeatedly.

Recent microengineering approaches have begun to treat curvature as an active and tunable mechanical cue. Controlled epithelial deformation has shown that curvature orientation modifies the balance between tissue tension and torque, with consequences for calcium signaling and gene expression (7). A hydraulically actuated organ-on-chip system has likewise generated millimeter-scale dome geometries in corneal stromal cultures, where geometric deformation promoted phenotypic and extracellular-matrix remodeling (4). Together, these advances establish the biological relevance of curvature while highlighting the need for platforms that combine bidirectional geometric control, repeated actuation, and real-time quantitative imaging.

A central unresolved question is therefore how epithelia respond when curvature changes over time, particularly when the same tissue alternates between geometries of opposite orientation. Resolving this question requires experimental control over deformation amplitude, rate, persistence, and history, together with three-dimensional measurements capable of separating cell and nuclear responses along the curved surface.

To address this need, we developed CurvoChip, a compact pneumatically driven microsystem for on-demand modulation of epithelial curvature. Building on micropneumatic strategies developed for cell mechanobiology (5), CurvoChip uses controlled pressure actuation of an ultrathin flexible membrane to generate reproducible concave and convex out-of-plane geometries. The platform provides control over deformation amplitude and temporal sequence, is compatible with time-lapse microscopy, and supports long-term culture under standard incubator conditions.

Here, we describe the design, fabrication, and mechanical calibration of CurvoChip and demonstrate its use with confluent MDCK epithelial monolayers. By combining live confocal microscopy with three-dimensional surface reconstruction and segmentation, we quantify how opposite curvature orientations remodel cell and nuclear architecture. Our results identify a pronounced asymmetry between concave and convex deformation and establish CurvoChip as a platform for investigating how epithelial tissues integrate dynamic geometry.

## 2. Materials and Methods

### 2.1 Principle of the dynamic curvature microsystem

CurvoChip is a pneumatically actuated device that applies concave, convex, or alternating deformations to epithelial monolayers cultured on a thin, flexible polydimethylsiloxane (PDMS) membrane. The microsystem is assembled from machined metal components (**Fig. 1A**) and supports both cell culture and live observation during controlled deformation (**Fig. 1B**). A programmable pressure controller (Cobalt, Elveflow) applies pressure beneath a 20-µm PDMS membrane bonded to a perforated thermoplastic disc. Negative and positive pressures deform the suspended membrane in opposite directions, generating concave cavities or convex domes relative to the epithelial apical surface. The system can be used in an open configuration inside a standard incubator or in a sealed configuration for on-stage imaging (**Fig. 1C**).

**Figure 1.**
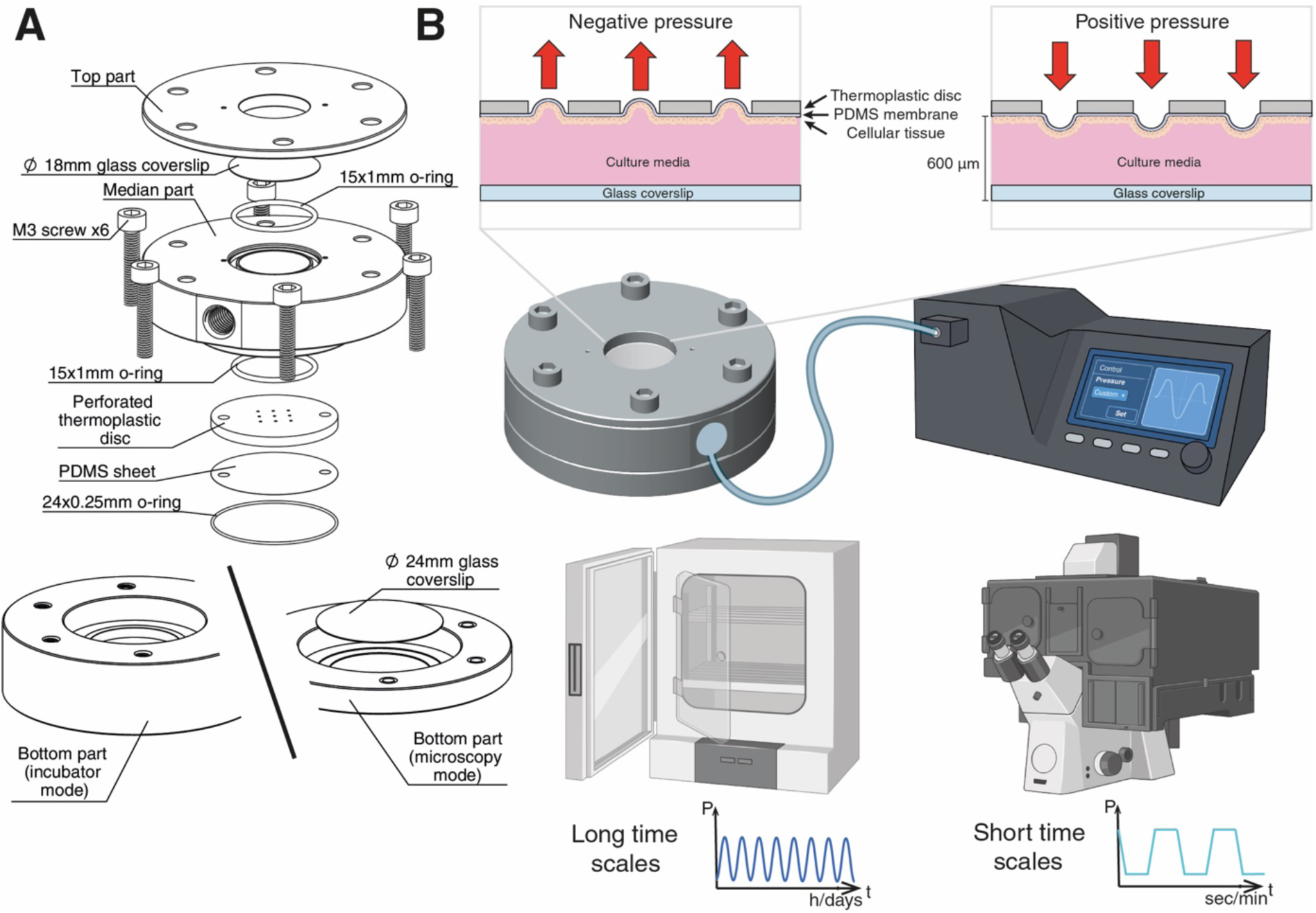
Concept of the CurvoChip microsystem. **(A)** Exploded view showing the stainless-steel housing, thermoplastic-PDMS disc, glass coverslips, and silicone seals. **(B)** Principle of curvature modulation and overview of the three-part housing forming the pneumatic and culture chambers. Negative and positive pressures deform the suspended membrane in opposite directions, generating concave or convex epithelial geometries. **(C)** Open incubator and sealed microscope configurations for long-term culture and real-time imaging, respectively.

### 2.2 Microsystem design and assembly

The CurvoChip housing comprises three stainless-steel plates that define a pneumatic chamber connected through Luer-lock fittings and a culture chamber containing the cell-seeded PDMS membrane (**Fig. 1A**). Two circular glass coverslips (18 and 25 mm in diameter) provide optical access. The 25-mm coverslip is separated from the culture surface by a 200-µm spacer, allowing imaging with 4× to 40× objectives having a working distance of at least 600 µm (**Fig. 1B**). Silicone O-rings provide leak-tight sealing, and lateral ports permit medium exchange. Stainless-steel parts were autoclaved at 121 °C for 20 min; the remaining components were sterilized with 70% ethanol and ultraviolet irradiation before assembly (**Fig. 2A 1)**. The functionalized thermoplastic-PDMS disc (**Fig. 2A 2)** was seeded at high density to establish a confluent epithelial monolayer (**Fig. 2A 3)** and then mounted aseptically in the device (**Fig. 2A 4)**. The final assembly was used either open for incubator experiments (**Fig. 2B 1-B2)** or sealed for microscopy (**Fig. 2C 1-C2)**.

**Figure 2.**
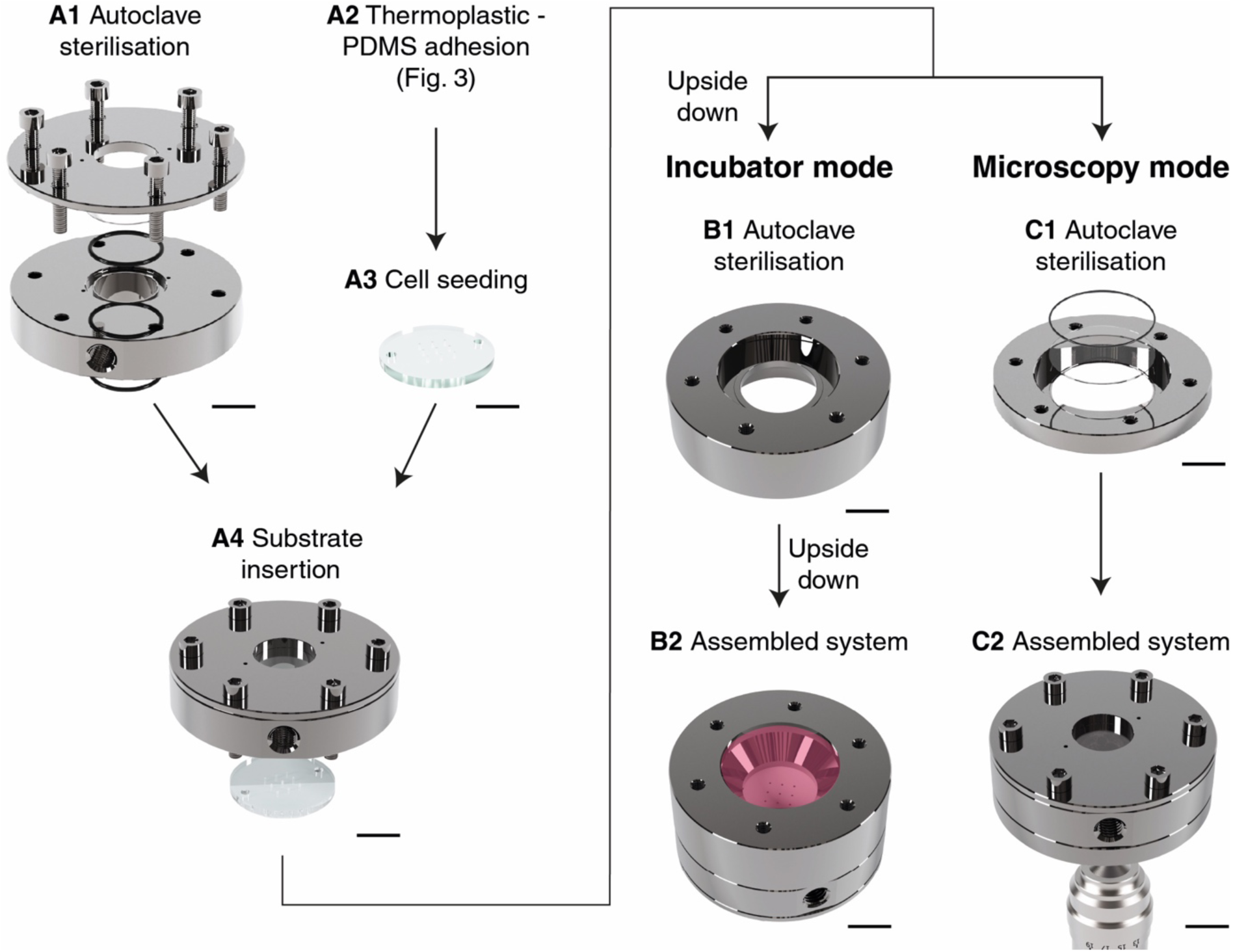
Assembly and operation of CurvoChip. **(A1-A4)** Sequential preparation and insertion of the thermoplastic-PDMS disc carrying the epithelial monolayer. **(B1-B2)** Open configuration for culture and actuation in a standard incubator. **(C1-C2)** Sealed configuration for on-stage live imaging. Scale bars, 1 cm.

### 2.3 Thermoplastic-PDMS bonding

Transparent polycarbonate (PC) or poly(methyl methacrylate) (PMMA) discs (25 mm diameter, 2 mm thickness; Goodfellow) were drilled with up to 49 circular apertures of 500 µm diameter, with a center-to-center spacing equal to twice the aperture diameter. Commercial 20-µm PDMS sheets (Limitless Shielding) were washed sequentially with isopropanol and deionized water and dried under nitrogen. PC discs were activated by O₂ plasma (29.6 W, 3 min; Harrick Plasma PDC-002) and incubated in 50% (v/v) APTES in water for 20 min (**Fig. 3A**). PMMA discs were spin-coated (2,000 rpm, 60 s) with a TEOS/chloroform/ethanol mixture (20:10:60, v/v/v) supplemented with 10% 0.1 M HCl, baked at 80 °C for 1 h, and activated by O₂ plasma (**Fig. 3B**). PDMS membranes were plasma-activated, brought into contact with the treated discs, and cured at 60 °C for 30 min under light pressure, yielding suspended deformable membranes across the apertures.

**Figure 3.**
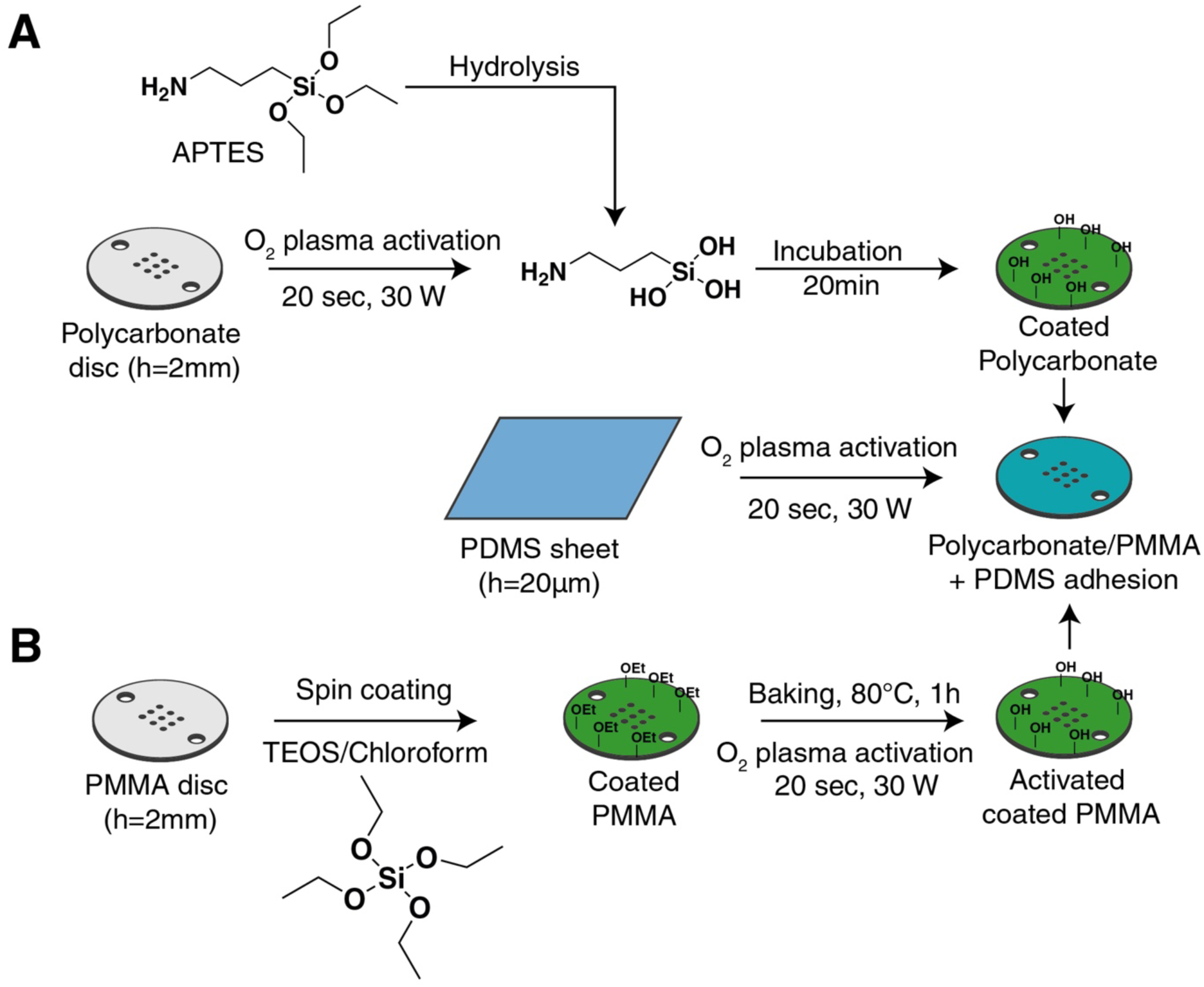
Surface functionalization and bonding procedures for thermoplastic–PDMS interfaces. **(A)** PDMS-PC bonding. Polycarbonate discs (2 mm thick) were activated by O₂ plasma and incubated for 20 min in 50% (v/v) APTES solution to introduce aminosilane groups. After hydrolysis, silanol-terminated APTES molecules formed an amine-functional coating on the PC surface. A 20-µm O₂-plasma-treated PDMS membrane was brought into contact with the functionalized PC and cured at 60 °C for 30 min to yield a strong covalent Si–O–Si linkage. **(B)** PDMS-PMMA bonding. PMMA discs (2 mm thick) were spin-coated with a TEOS/chloroform solution to deposit a thin silicate layer, aged overnight, and plasma-activated to generate surface hydroxyl groups. The activated PMMA surface was then bonded to an O₂-plasma-treated PDMS membrane, forming a permanent PMMA–PDMS hybrid suitable for pneumatic actuation.

### 2.4 Membrane profilometry and finite-element modeling

Membrane deformation was quantified from confocal z-stacks of fluorescently labeled PDMS membranes acquired during stepwise pressure actuation. Surface profiles and the central deflection (δ) were extracted for each aperture and compared with the analytical scaling relations described in the Supplementary Theory. Finite-element simulations were performed in COMSOL Multiphysics using a Neo-Hookean constitutive model with a PDMS Young’s modulus of 6 MPa. The membrane was fixed at the aperture boundary and subjected to a spatially uniform normal pressure. Simulated profiles and central deflections were exported over the experimental pressure range and compared directly with confocal measurements.

### 2.5 Cell culture

MDCK cells stably expressing E-cadherin-mCherry were maintained in Dulbecco’s modified Eagle medium supplemented with 10% fetal bovine serum and 1% penicillin-streptomycin. Before seeding, PDMS surfaces were coated with fibronectin (25 µg mL⁻¹, 1 h, 37 °C). Cells were seeded at 3 × 10⁵ cells per disc and cultured to confluence. For live-cell imaging of nuclear morphology, the culture medium was replaced with FluoroBrite DMEM and nuclei were labeled with Hoechst 33342 (0.5 µL mL⁻¹, 1 h).

### 2.6 Live confocal microscopy

Live imaging was performed on a Nikon Ti2-A1R laser-scanning confocal microscope equipped with a Plan Apo 20× air objective (NA 0.75) and an environmental chamber maintained at 37 °C and 5% C0_2_. Three-dimensional stacks spanning the full height of the deformed monolayer were acquired with a 1-µm z-step. For acute deformation experiments, pressure was ramped from 0 to ±400 mbar at 1 mbar s⁻¹, and z-stacks were acquired within 15 min after completion of the ramp. E-cadherin-mCherry and Hoechst fluorescence were used to visualize cell-cell junctions and nuclei, respectively. Images were acquired and initially processed in NIS-Elements AR (Nikon).

### 2.7 Epithelial surface reconstruction and morphometric analysis

Fluorescence signals corresponding to cell–cell junctions (E-cadherin) were used for three-dimensional reconstruction and segmentation of epithelial tissues. E-cadherin image stacks were denoised, intensity-normalized, and resampled to isotropic voxels before analysis in MorphoGraphX v2 (8). The epithelial surface was extracted in three dimensions, represented as a triangular mesh, and used to project the junctional fluorescence signal onto the local apical surface. Cell centers were seeded on the projected signal, and individual cells were segmented using a watershed-based algorithm. Quantitative morphometric parameters, including cell area, perimeter, and spatial coordinates were extracted from the segmented surface and processed with custom Python scripts. To resolve spatial variation, cells were assigned to concentric zones from the flat region (Zflat) toward the dome or cavity apex (Z1-Z3). Cells at the aperture boundary, where curvature reverses locally and mechanical constraints differ, were excluded. For each condition, measurements were pooled from multiple independent domes and imaging fields, and dentical processing parameters were used across conditions to ensure consistency and reproducibility.

### 2.8 Nuclear segmentation and morphometry

Hoechst-labeled nuclei were segmented in three dimensions from confocal stacks using intensity thresholding followed by watershed separation of touching objects. Segmented objects intersecting image boundaries or falling outside the epithelial surface mask were excluded. For each nucleus, the projected area in the local epithelial plane, thickness along the local surface normal, and three-dimensional volume were quantified. Nuclear measurements were assigned to the same Zflat, Z2, and Z3 regions used for cellular morphometry.

### 2.9 Statistical analysis

Data are shown as distributions of individual cells together with independent-sample means and are summarized as mean ± SD. Statistical comparisons were performed in GraphPad Prism using the Kruskal-Wallis test followed by Dunn’s multiple-comparisons test. A two-sided p value < 0.05 was considered significant (*p < 0.05; **p < 0.01; ***p < 0.001; ****p < 0.0001).

## 3. Results

To investigate how dynamic curvature modulations influence epithelial monolayer behavior, we first focused on concave and convex spherical deformations generated by actuating the membrane through circular apertures in the rigid thermoplastic support. When an elastic membrane is pneumatically actuated through circular openings, the suspended membrane formed a cup-like (concave) or dome-like (convex) surface relative to the epithelial apical side, depending on pressure direction In both configurations, the surface remains locally spherical, with the two principal curvatures being equal in magnitude. Consequently, the Gaussian curvature is positive for both outward and inward hemispheres, as it depends on the product of the principal curvatures. In contrast, the mean curvature changes sign: it is positive for convex domes and negative for concave invaginations. While Gaussian curvature characterizes the intrinsic surface type (spherical versus saddle-like), mean curvature encodes the direction of bending, a distinction that is particularly relevant for biological systems, which may differentially sense curvature orientation and asymmetry.

Importantly, CurvoChip is not restricted to circular apertures or static hemispherical profiles. By changing aperture geometry and the temporal pressure program, the platform can in principle generate spherical, cylindrical, saddle-like, and multicurved landscapes with reversible temporal modulation **(Supplementary Fig. 1)**. This capability mirrors the conceptual framework of the global curvature patterns described previously (9) and establishes CurvoChip as a versatile experimental system to probe epithelial responses to complex and physiologically relevant curvature cues under dynamic conditions. On this basis, we first sought to quantitatively characterize the mechanical behavior of the deformable PDMS membrane.

### 3.1 Mechanical characterization and modeling of PDMS deformation

We derived analytical scaling relations to describe how aperture radius *(R)*, membrane thickness *(h)*, PDMS Young’s modulus *(E)*, and applied pressure *(P)* determine central deflection *(δ)* and radius of curvature *(ρ)* (**Fig. 4A**). In the stretching-dominated regime, the inflated membrane can be approximated as a pressurized spherical cap, for which Laplace’s law gives:

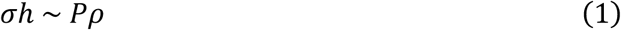

where *σ* is the membrane stress. Assuming linear elasticity at small strain, Hooke’s law gives (see Supplementary Theory):

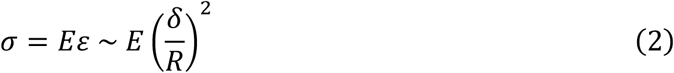

Combining expressions (1) and (2) yields:

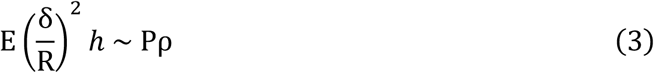

which leads to:

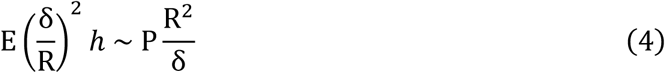

or equivalently:

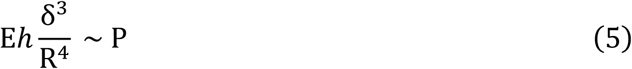

and thus the normalized deflection:

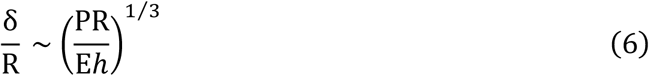

Thus, in the stretching-dominated regime, normalized deflection *δ/R* scales with the cubic root of *PR/Eh*. At small deflections or for comparatively thick membranes, bending is no longer negligible, and the spherical-cap approximation becomes less accurate. In this bending-dominated regime, the flexural moment *(M)* can be expressed as:

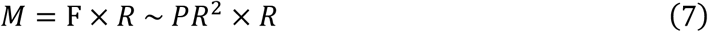

*F* being the force applied on the disc due to the pressure. Another expression of *M* is:

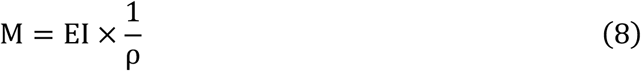

With *I* the quadratic moment:

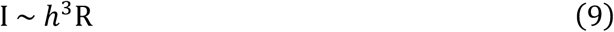

These relations yield:

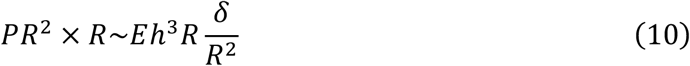

From which follows:

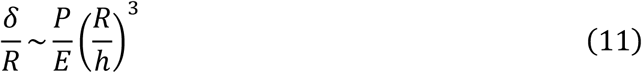

corresponding to a regime in which deflection scales approximately linearly with pressure. The deformed membrane approaches a spherical-cap profile when the boundary-layer length l (**Fig. 4A**) satisfies:

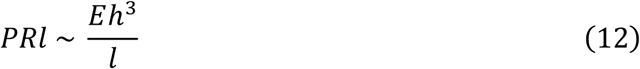

which gives:

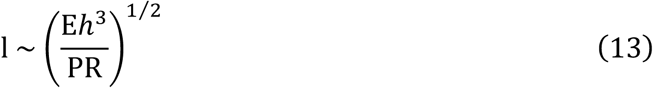

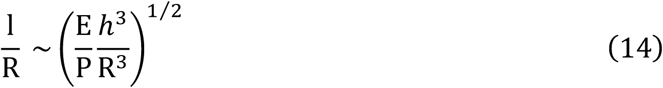

The profile is therefore approximately hemispherical when l/R is negligible:

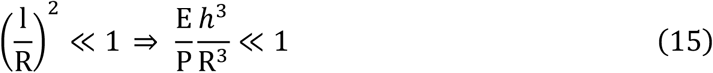

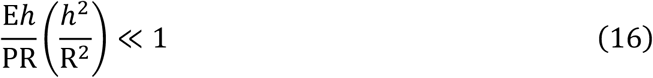

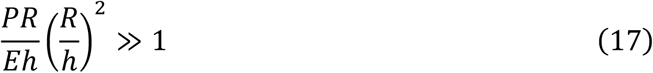

This condition is favored by high pressure, a large aperture radius, a low Young’s modulus, and a thin membrane.

We tested these predictions by confocal profilometry of fluorescently labeled PDMS membranes (**Fig. 4B**) and finite-element simulations using a Neo-Hookean material model (**Fig. 4C**). The analytical scaling captures the dominant low-to-intermediate pressure trend but increasingly underestimates deflection at the largest loads, where finite strain, clamped-boundary effects, and deviations from the idealized spherical-cap assumptions become important. By contrast, simulated profiles reproduced the experimental measurements within approximately 10% across the tested range (**Fig. 4D**). The finite-element model was therefore used as the quantitative calibration framework for selecting CurvoChip pressure programs.

**Figure 4.**
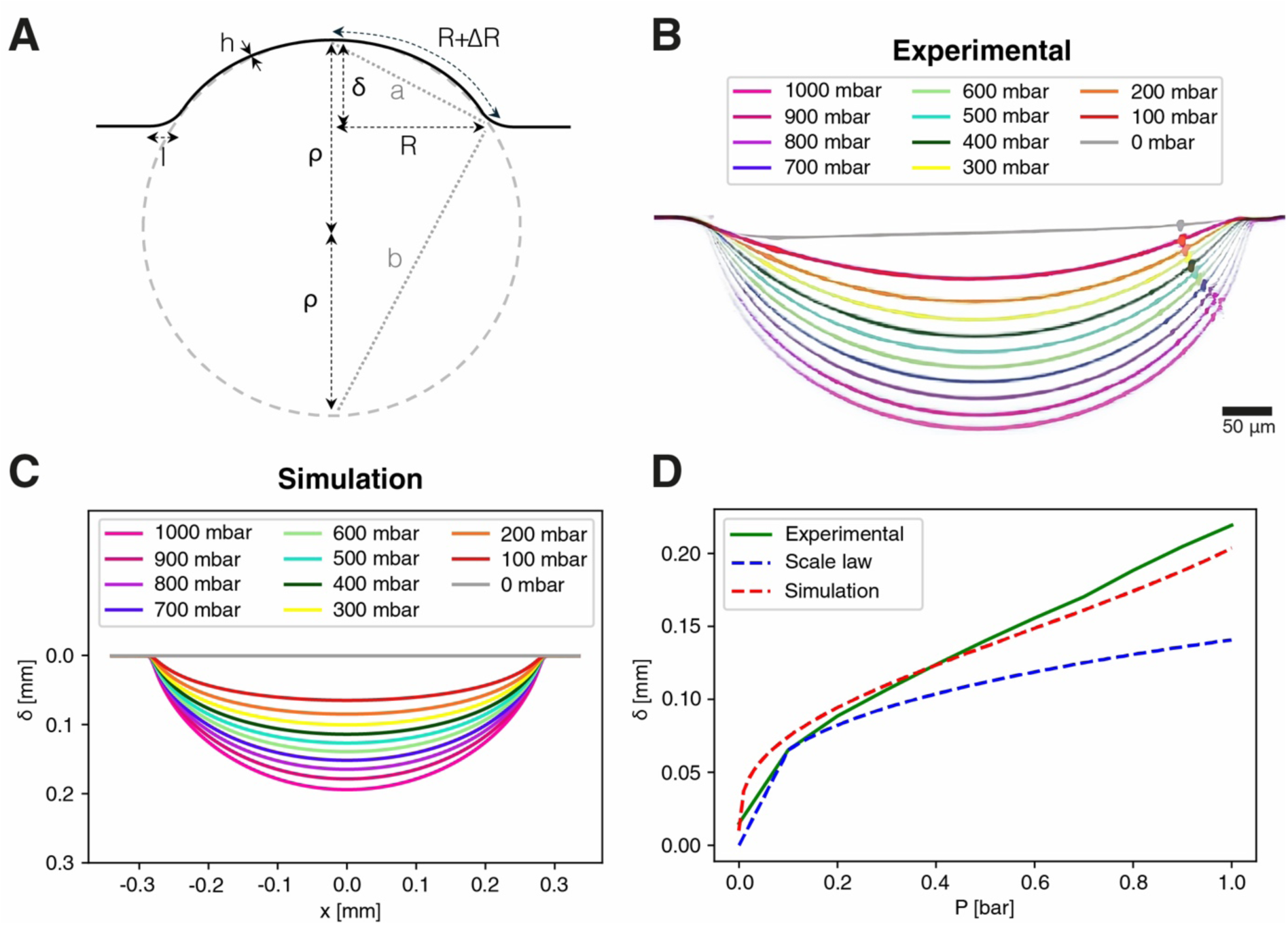
Modeling and experimental characterization of PDMS deformation. **(A)** Geometric parameters used in the analytical scaling analysis. **(B)** Experimental confocal profiles of fluorescently labeled PDMS membranes from 0 to 1,000 mbar. **(C)** Corresponding finite-element profiles generated with a Neo-Hookean material model. **(D)** Comparison of measured central deflection with analytical scaling and finite-element simulation. Scale bars, 100 µm.

### 3.2 Reversibility and stability under cyclic actuation

To assess long-term mechanical stability, PDMS membranes were subjected to 120 cycles alternating between −400 and +400 mbar over approximately 4 h. Each loading phase lasted 110 s and was followed by 10 s at baseline pressure (**Fig. 5A**).

**Figure 5.**
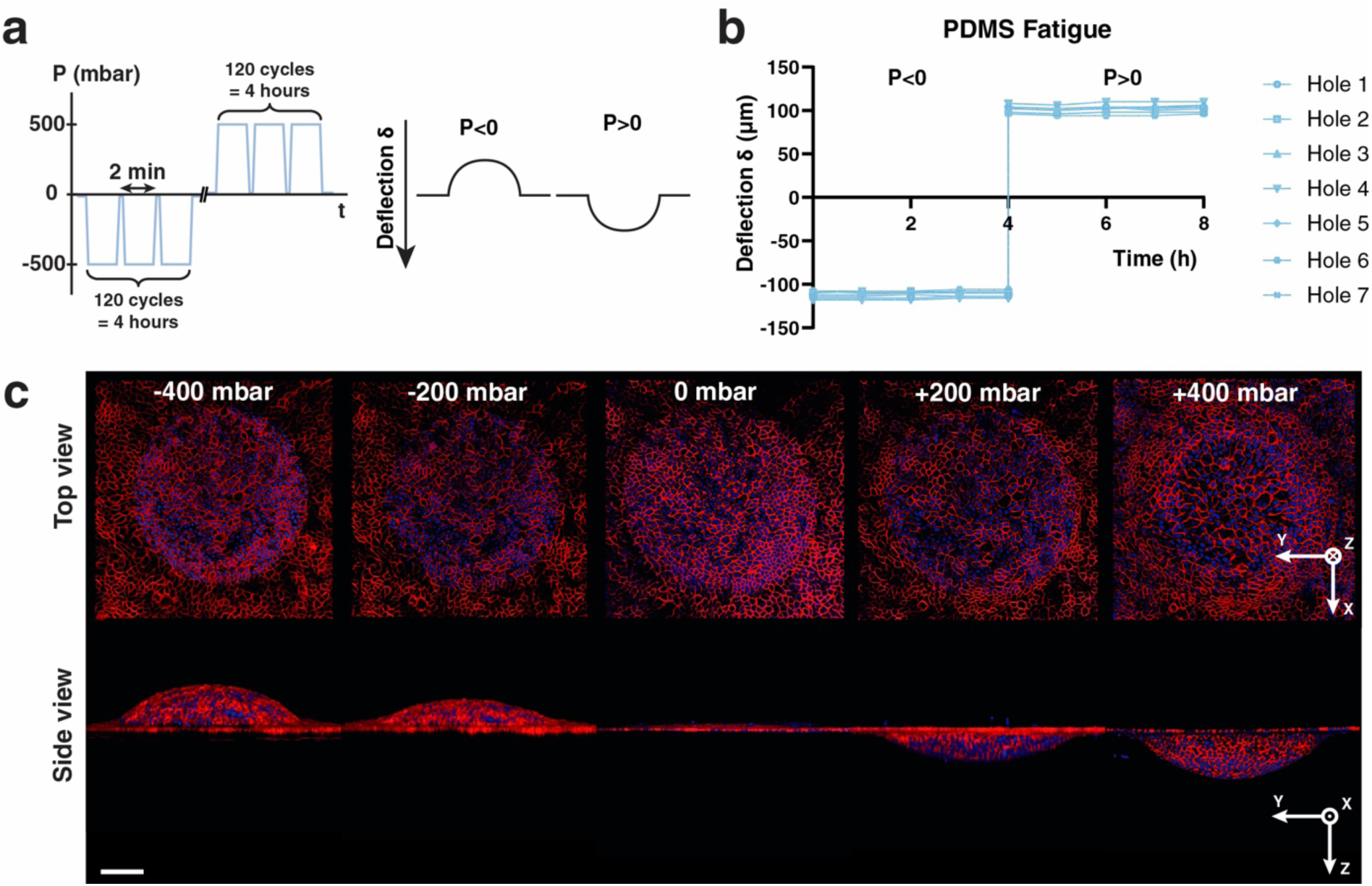
Fatigue analysis under cyclic curvature modulation. **(A)** Pressure programme alternating between −400 and +400 mbar, with 110-s loading phases and 10-s returns to baseline. **(B)** Central membrane deflection measured hourly at seven apertures over 120 cycles. (C) Representative confocal cross-sections at −400, −200, 0, +200, and +400 mbar. Scale bar, 200 µm.

Confocal z-profiles of fluorescently marked PDMS membranes were recorded hourly at seven circular apertures to quantify central deflection and potential drift. The mean deflection amplitude remained stable at 105 ± 2 µm throughout the experiment (**Fig. 5B).** No measurable hysteresis or residual deformation was observed between consecutive inflation and deflation phases, confirming the excellent elastic recovery of the thin PDMS sheet and the reliability of the thermoplastic–PDMS bonding interface. Representative cross-sections confirmed reproducible membrane shapes across negative, zero, and positive pressures (**Fig. 5C**).

These measurements demonstrate that the thermoplastic-PDMS interface and suspended membrane withstand repeated bidirectional actuation over the duration required for live-cell experiments. CurvoChip can therefore deliver reproducible curvature histories without measurable mechanical fatigue over at least 120 cycles.

### 3.3 Three-dimensional morphometric analysis on curved epithelia

To quantify epithelial architecture directly on the deformed surface, we established a semi-automated analysis workflow combining MorphoGraphX v2 (8) with custom Python scripts (**Fig. 6**). The pipeline extracts morphological descriptors, including cell and nuclear area, perimeter, orientation, and shape factor from both curved regions (domes or cavities) and adjacent flat controls while retaining each cell’s radial position.

**Figure 6.**
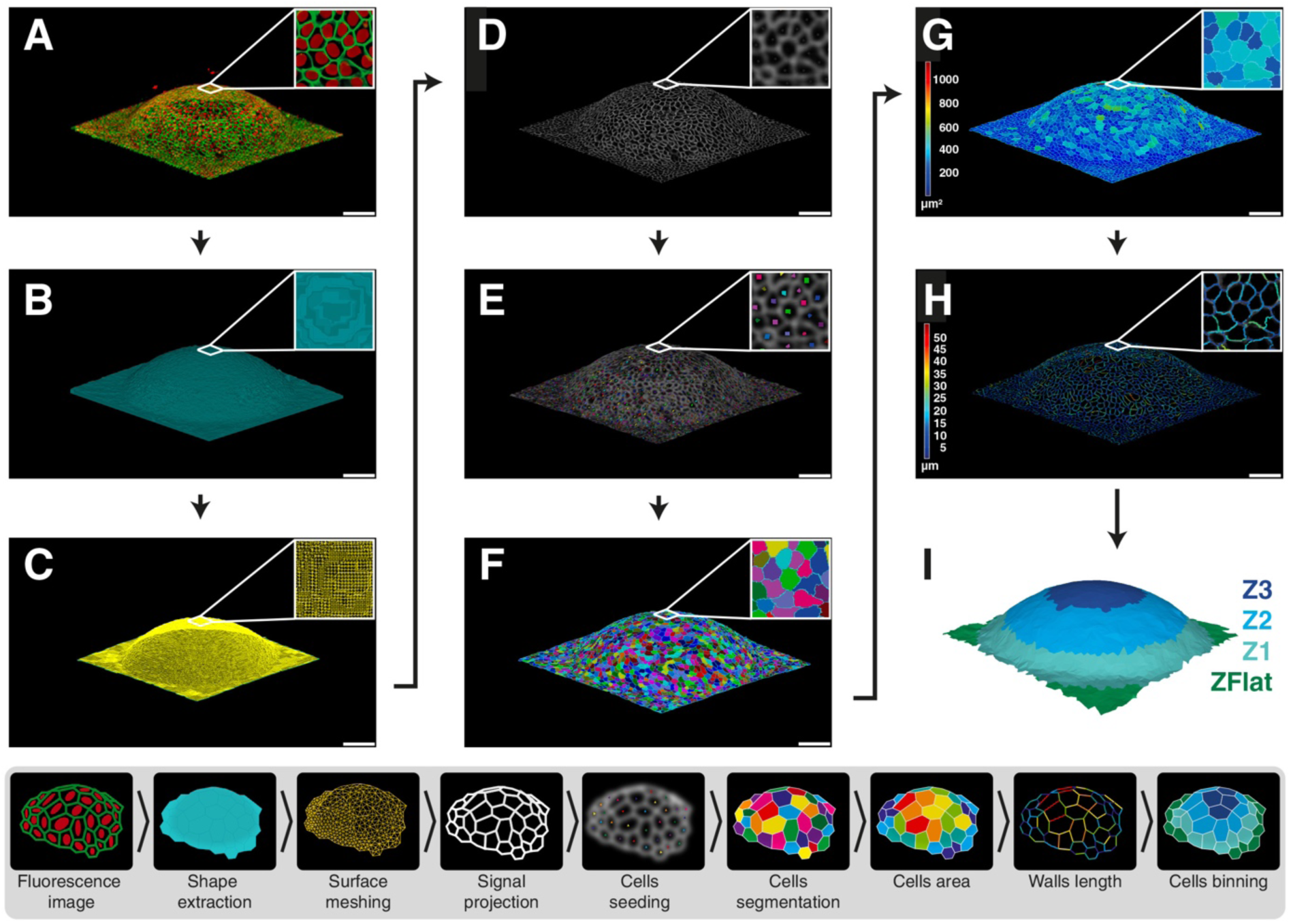
Three-dimensional 3D morphometric pipeline for curvature analysis. **(A)** Raw three-dimensional E-cadherin-mCherry fluorescence. **(B)** Epithelial surface extraction. **(C)** Surface meshing. **(D)** Projection of the junctional signal onto the local epithelial surface. **(E)** Cell-center seeding. **(F)** Watershed segmentation of individual cells. **(G)** Cell-area map. **(H)** Cell-wall-length map. **(I)** Assignment of cells to concentric regions from the flat periphery (Zflat) to the curved apex (Z1-Z3). Scale bars, 50 µm.

Raw confocal stacks of E-cadherin-mCherry monolayers were denoised and resampled to isotropic voxels (**Fig. 6A**). The epithelial surface was extracted (**Fig. 6B**), fitted with a smooth reference mesh following the PDMS deformation (**Fig. 6C**), and used to project junctional fluorescence onto onto a two-dimensional map corresponding to the local apical plane (**Fig. 6D**). The projected signal was subsequently seeded in the center of the cells (**Fig. 6E**) and individual cell boundaries were segmented by watershed (**Fig. 6F**). From these segmented masks, single-cell geometrical features such as cell area (**Fig. 6G**), perimeter (**Fig. 6H**), and the position of the elements composing the meshed surface were extracted. Morphometric data can be organized into concentric radial zones extending from the dome apex (Z1, central 2/5^th^ of the dome) to the peripheral flat region (Z3, most peripheral 1/5^th^ of the dome), with an additional flat control zone (Zflat) outside the deformed area (**Fig. 6I**).

This workflow provides robust and reproducible single-cell measurements across non-planar epithelia and enables direct comparison of spatially matched regions under opposite curvature orientations.

### 3.4 Opposite curvature orientations asymmetrically remodel epithelial architecture

To investigate how epithelial cells respond to dynamic curvature modulation, confluent MDCK monolayers were subjected to acute bidirectional deformation by ramping pressure from 0 to −400 or +400 mbar at 1 mbar s⁻¹ (**Fig. 7A**).

**Figure 7.**
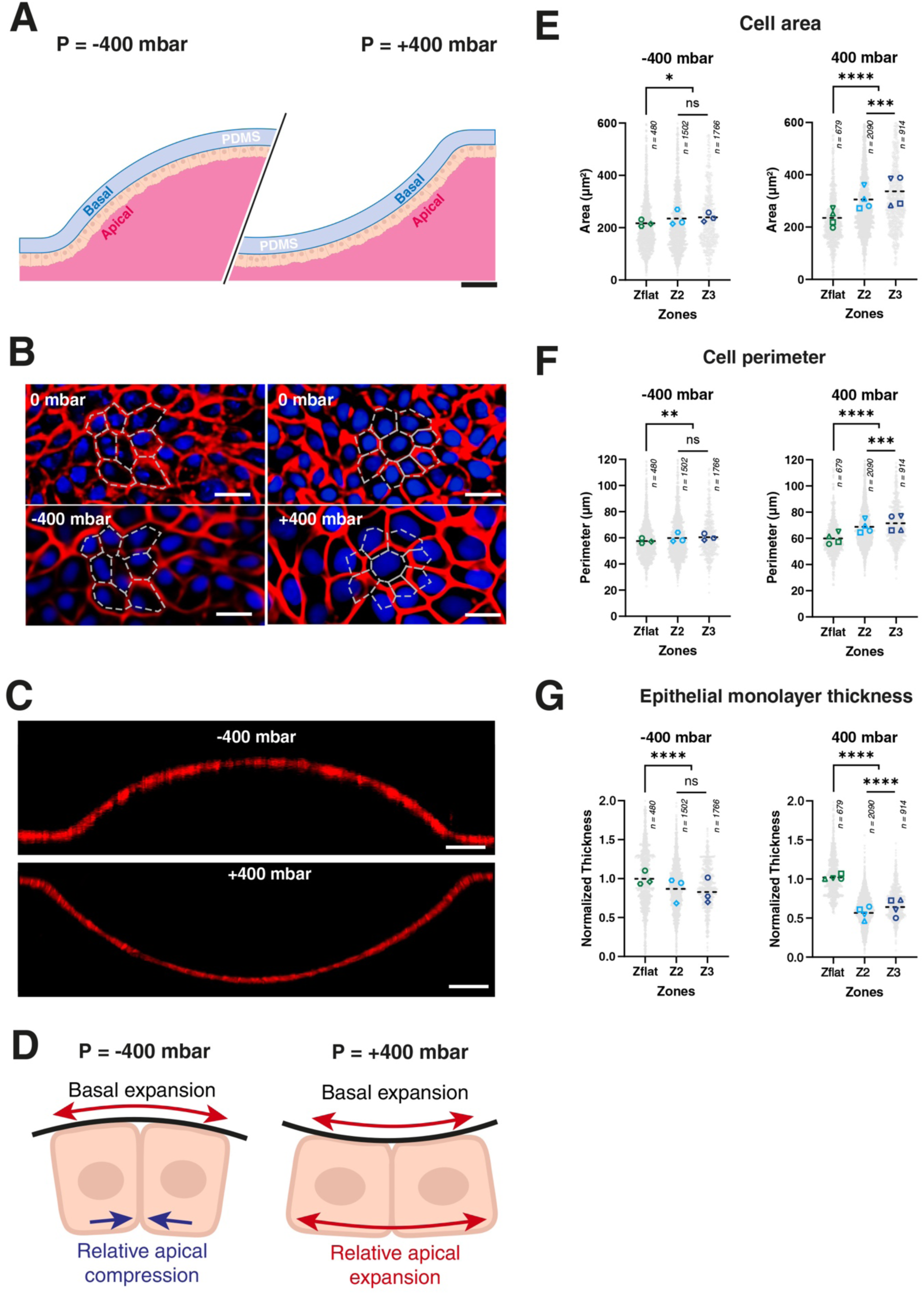
Curvature-orientation-dependent remodeling of epithelial architecture. **(A)** MDCK monolayers under concave (−400 mbar) and convex (+400 mbar) deformation. **(B)** Apical views before and after deformation; E-cadherin-mCherry is shown in red and nuclei in blue. **(C)** Cross-sectional views of deformed monolayers. Pressure was ramped from 0 to ±400 mbar over 6 min 40 s (1 mbar s⁻¹). **(D)** Schematic of the relative apical and basal geometric changes imposed by the two orientations. Cell area **(E)**, cell perimeter **(F)**, and normalized epithelial thickness **(G)** in flat (Zflat) and curved (Z2 and Z3) regions. Grey points represent individual cells; colored symbols denote independent samples. Scale bars, 50 µm (A,C) and 20 µm (B). Statistical significance is defined in Methods.

Live confocal imaging of E-cadherin-mCherry junctions and Hoechst-labeled nuclei showed that concave deformation (cup) produced comparatively modest changes in apical cell shape, whereas convex deformation (dome) caused a pronounced increase in apical cell area, particularly near the apex. (**Fig. 7B**). Interestingly, cross-sectional reconstructions further suggested epithelial thinning under both orientations, with a stronger effect in convex regions (**Fig. 7C**).

To verify that curvature modulates not only the in-plane cell geometry but also the vertical organization of the epithelial monolayer, we used the image-analysis pipeline to quantify cell area (**Fig. 7D**) and perimeter (**Fig. 7E**) in the regions near the apex (zones Z2 and Z3) and in adjacent flat control areas, as well as to measure the monolayer thickness in these regions. Our analysis revealed a slight but significant increase in cell area within concave regions (238 ± 17 µm^2^) compared with flat zones (216/234 ± 16/33 µm^2^), whereas convex curvature induced a much larger increase in cell spreading (336 ± 58 µm^2^) (**Fig. 7E**). Notably, curvature-induced changes in cell area were comparable across concave regions, with no significant differences observed between zones Z2 and Z3. In contrast, within convex regions, cells located at the apex (Z3) exhibited a significantly larger area than those in Z2 and in flat regions. Together, these results reveal a spatially homogeneous cellular response along concave surfaces, whereas convex curvature induces a graded modulation of cell area. Interestingly, similar trends were observed for cell perimeter, which was slightly but significantly higher in concave regions (61 ±3 µm) relative to flat controls (57/60 ±2/4 µm), and substantially larger in convex regions (72 ±6 µm) (**Fig. 7F**). Finally, the epithelial monolayer thickness was quantified from high-resolution confocal side-view profiles (**Fig. 7G**). The results showed a reduction in epithelial thickness in both concave and convex zones. However, the decrease in thickness of zones Z2 and Z3, normalized to flat regions (**Fig. 7G**), was significantly greater in convex curvature zones (0.67 ± 0.11) than in concave curvature zones (0.84 ± 0.16).

Altogether, these results indicate that epithelial morphology depends on curvature orientation as well as magnitude. Convex deformation promotes pronounced apical expansion and thinning, whereas concave deformation elicits a weaker and more spatially uniform response, consistent with distinct apico-basal geometric constraints in the two configurations.

### 3.5 Curvature orientation differentially reshapes epithelial nuclei

We next asked whether the curvature-orientation-dependent cellular response extended to the nucleus, a mechanically integrated organelle whose shape reflects both cytoskeletal forces and spatial constraints (**Fig. 8A**). Hoechst-labeled nuclei in epithelial monolayers subjected to acute deformation from 0 to either −400 or +400 mbar at 1 mbar s⁻¹ (**Fig. 8B**) were segmented in three dimensions (**Fig. 8C**), enabling measurement of nuclear projected area, thickness, and volume (**Fig. 8A**) in the flat, Z2, and Z3 regions.

**Figure 8.**
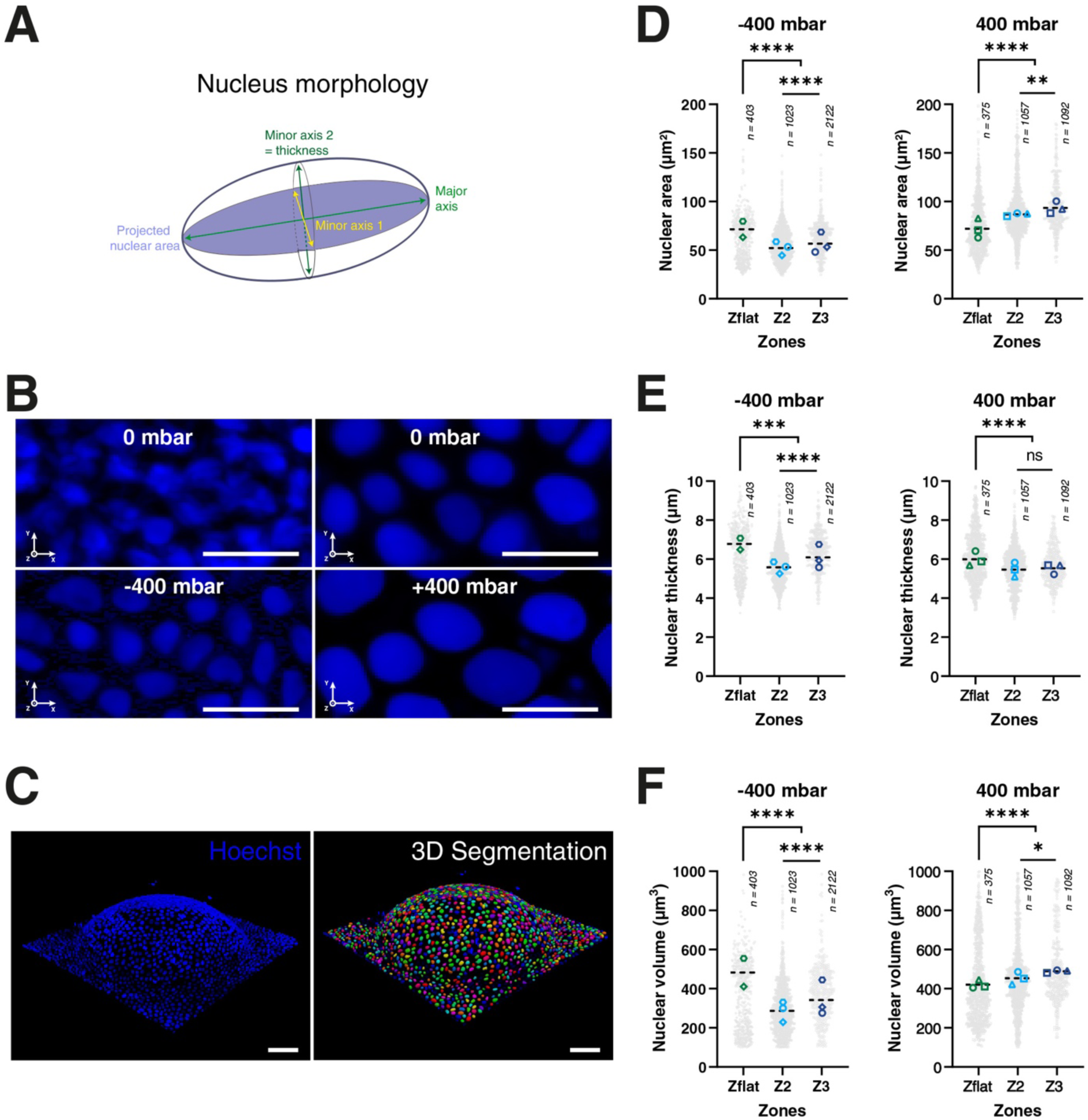
Curvature-orientation-dependent remodeling of epithelial nuclei. **(A)** Nuclear morphometric descriptors. **(B)** Apical views of Hoechst-labeled nuclei before and after concave (−400 mbar) or convex (+400 mbar) deformation. **(C)** Three-dimensional Hoechst signal and segmented nuclei on a representative curved monolayer. Nuclear projected area **(D)**, thickness **(E)**, and volume **(F)** in flat (Zflat) and curved (Z2 and Z3) regions. Grey points represent individual nuclei; colored symbols denote independent samples. Scale bars, 20 µm (B) and 50 µm (C). Statistical significance is defined in Methods.

Under concave deformation, nuclear projected area decreased sharply in Z2 and partially recovered in Z3. Conversely, convex deformation caused a progressive increase in projected nuclear area from Zflat to Z2 and Z3 (**Fig. 8D**). Nuclear thickness also varied with curvature orientation and position. Under concave deformation, it decreased in Z2 and partially recovered in Z3, whereas under convex deformation, it decreased in Z2 and remained similar between Z2 and Z3 (**Fig. 8E**). Nuclear volume exhibited a comparable orientation-dependent response: concave deformation produced a marked reduction in Z2 followed by partial recovery in Z3, while convex deformation induced a modest but progressive increase across the two curved regions (**Fig. 8F**).

Together, these results demonstrate that opposite curvature orientations induce distinct three-dimensional nuclear remodeling. Convex deformation combines lateral nuclear expansion with reduced thickness and a modest increase in volume, whereas concave deformation reduces nuclear projected area, thickness, and volume, with the strongest effects observed in Z2. These nuclear responses mirror the asymmetric remodeling of epithelial cell and tissue architecture, demonstrating that dynamic geometry propagates from the tissue scale to intracellular structures.

## 4. Discussion and outlook

CurvoChip provides reversible control over epithelial geometry while retaining optical access, standard culture conditions, and compatibility with long-term live imaging. The combination of a reusable metal housing, exchangeable thermoplastic-PDMS inserts, and programmable pneumatic actuation makes it possible to prescribe deformation direction, amplitude, rate, and sequence. Its mechanical calibration and stability over 120 bidirectional cycles establish a practical operating window for experiments that move beyond static curved substrates toward explicitly time-dependent geometric histories.

The analytical treatment identifies the bending- and stretching-dominated regimes that govern membrane deflection and provides useful design rules for selecting aperture size and membrane thickness. Its divergence from experiment at the highest pressures is expected because the scaling analysis neglects finite-strain hyperelasticity, detailed clamped-edge mechanics, and departures from an ideal spherical cap. The close agreement of the Neo-Hookean finite-element model with measured profiles provides a more accurate calibration at large deformation and can be extended to non-circular apertures. The simulated cylindrical, saddle-like, and multicurved geometries illustrate how CurvoChip could be used to navigate a broader mean-curvature/Gaussian-curvature space than the spherical configurations tested here.

Our epithelial measurements reveal a pronounced response to curvature orientation. At matched pressure magnitude, convex deformation induced stronger cell spreading and tissue thinning than concave deformation, and the effect increased toward the apex. This asymmetry is consistent with the fact that opposite bending orientations impose different relative changes on the apical and basal surfaces of a polarized epithelium. The associated nuclear response reinforces this interpretation: convex deformation promoted lateral nuclear expansion and flattening, whereas concave deformation reduced nuclear projected area and volume. These distinct shape changes provide a direct route by which dynamic tissue geometry could influence chromatin organization, nuclear transport, and mechanosensitive transcription (10).

An important limitation is that pneumatic membrane deformation changes curvature together with in-plane strain, membrane tension, and pressure history. CurvoChip therefore measures the integrated epithelial response to a controlled geometric deformation rather than curvature in complete mechanical isolation. Comparing concave and convex states at matched pressure and similar deflection helps resolve the contribution of orientation, but future experiments should combine local strain mapping, force measurements, and pressure-only controls to separate these coupled variables. The current biological validation is also restricted to acute responses in MDCK monolayers; longer conditioning protocols, additional epithelial models, and molecular perturbations will be required to determine whether repeated curvature writes persistent mechanical states or modifies cell fate (11).

Future developments will extend the accessible actuation-frequency range, integrate perfusion for coordinated chemical and mechanical stimulation, and combine curvature control with traction-force microscopy, FRET-based tension sensors, and live reporters of ERK, YAP, calcium, or nuclear-envelope integrity. Parallelization across the patterned apertures should also enable systematic comparisons of curvature amplitude, duration, sequence, and recovery. These capabilities position CurvoChip as a versatile platform for studying morphogenesis, regeneration, barrier remodeling, and curvature-associated disease in dynamically evolving epithelial tissues.

## Acknowledgments

M.L. and S.G. acknowledge support from the University of Mons (UMONS), the FEDER Prostem Research Project no. 1510614 (Wallonia DGO6), the F.R.S.-FNRS Epiforce Project no. T.0092.21, the F.R.S.-FNRS Cellsqueezer Project no. J.0061.23, and the F.R.S.-FNRS Optopattern Project no. U.NO26.22. This work was supported by the Interreg projects ANTIRESI and MICROPLAITE, co-funded by the European Union through the European Regional Development Fund (ERDF-FEDER) under the Interreg France-Wallonie-Vlaanderen programme. M.L. is a Postdoctoral Researcher (Chargée de recherches) of the F.R.S.-FNRS. R.T. acknowledges doctoral support from the F.R.S.-FNRS (FRIA). S.G. acknowledges support from the Fonds pour la Recherche Médicale dans le Hainaut and the Francqui Foundation through a Francqui Research Professorship. The authors thank Gilles Simon (ILMTech platform) for assistance and discussions regarding CurvoChip fabrication.

## Author contributions

M.L. and S.G. conceptualized the study. R.T. performed the experiments and COMSOL simulations. All authors contributed to data interpretation, figure preparation, and manuscript writing and approved the final version.

## Competing interests

The authors declare no competing interests.

---

**Supplementary Figure 1.**
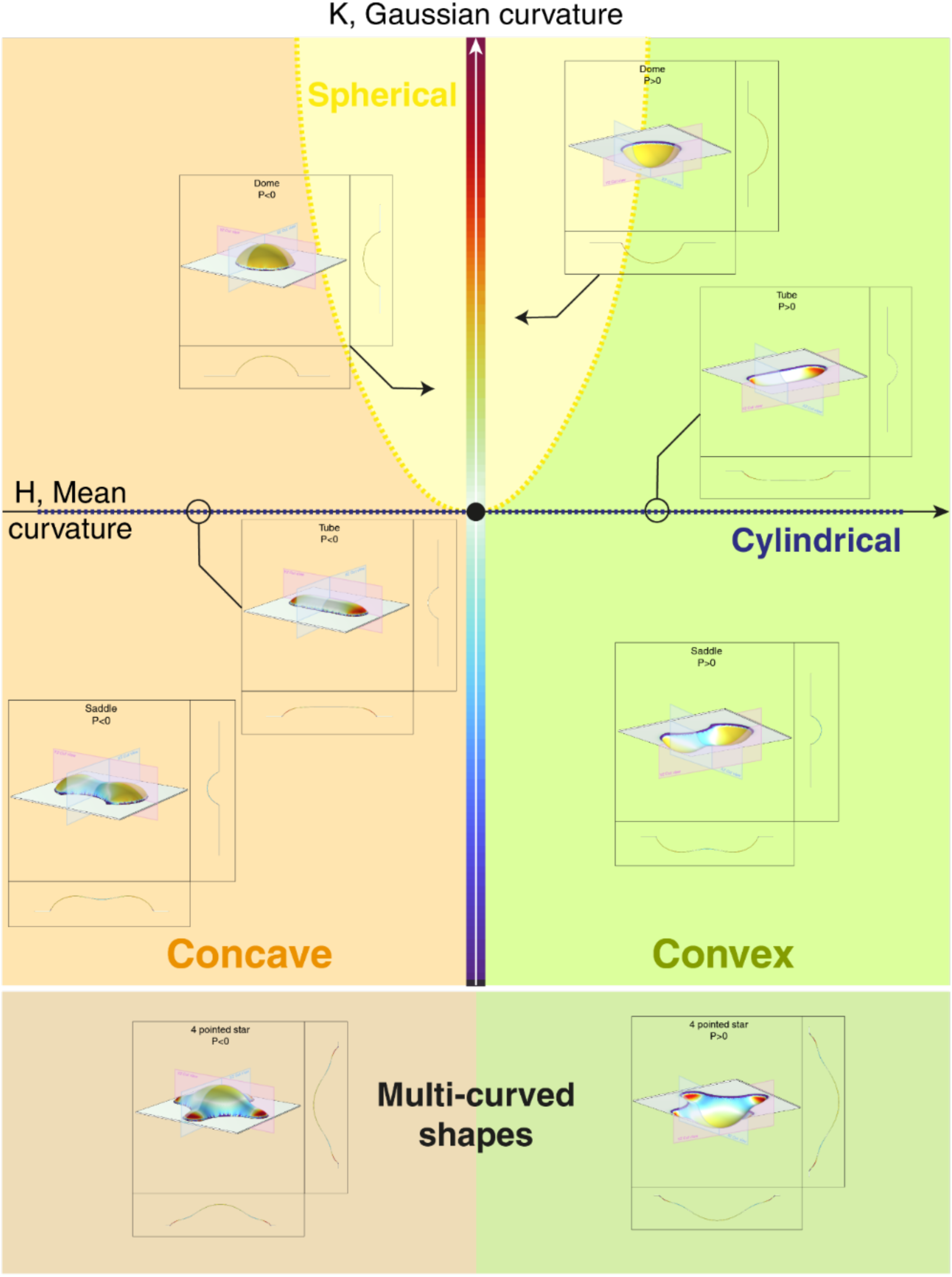
Numerical exploration of CurvoChip curvature landscapes. Finite-element simulations illustrate the range of profiles generated by changing the aperture geometry in the rigid thermoplastic support and the direction of pneumatic pressure. Representative spherical, cylindrical, saddle-like, and multicurved geometries are shown for negative (P < 0) and positive (P > 0) actuation, together with orthogonal cross-sections and three-dimensional reconstructions. The examples span positive, near-zero, and negative Gaussian curvature and both orientations of mean curvature, as summarized in the H-K curvature diagram.

## Supplementary Theory

Derivation supporting Equation (2)

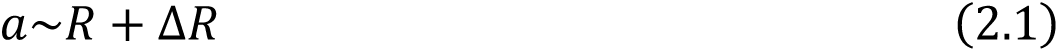

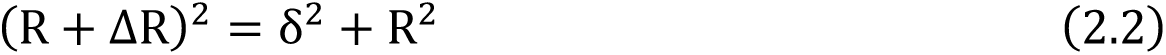

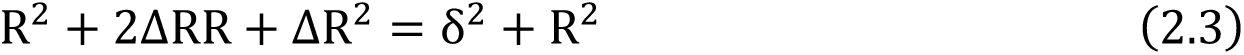

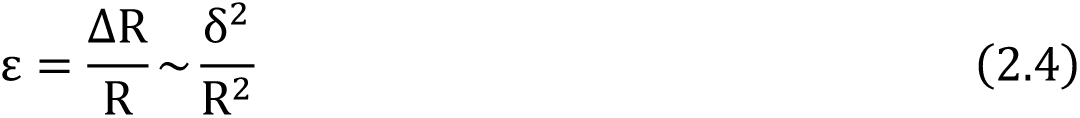

Derivation supporting Equation (4)

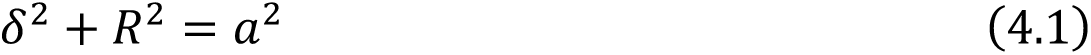

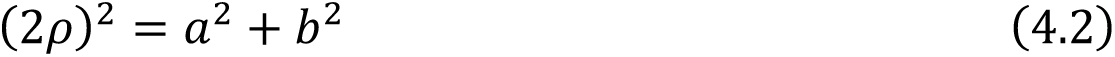

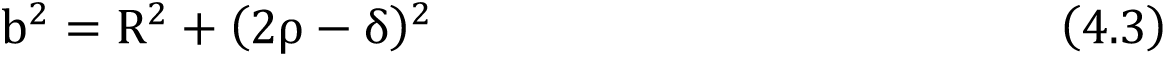

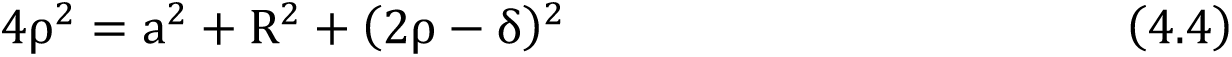

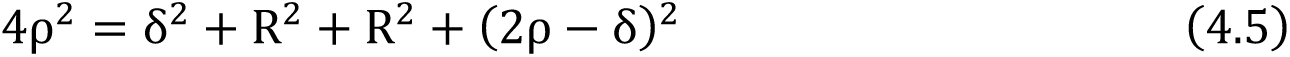

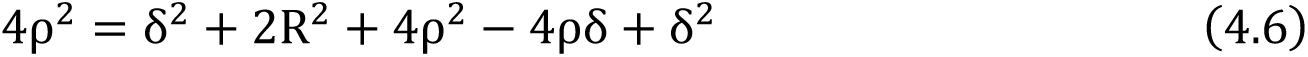

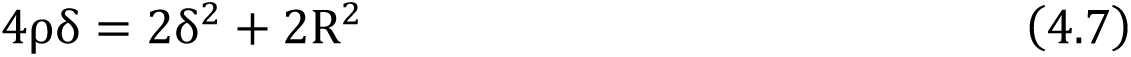

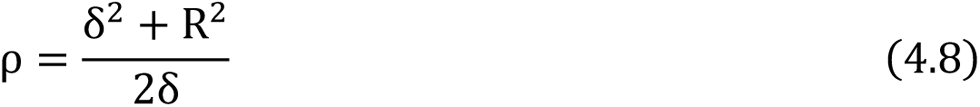

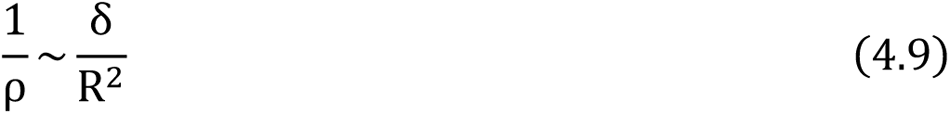

## Notes

### Competing Interest Statement

The authors have declared no competing interest.

